# Maturing neutrophils of lower density associate with thrombocytopenia in Puumala orthohantavirus-caused hemorrhagic fever with renal syndrome

**DOI:** 10.1101/2024.02.19.580937

**Authors:** Luz E. Cabrera, Johanna Tietäväinen, Suvi T. Jokiranta, Satu Mäkelä, Antti Vaheri, Jukka Mustonen, Olli Vapalahti, Mari Kanerva, Tomas Strandin

## Abstract

Puumala orthohantavirus-caused hemorrhagic fever with renal syndrome (PUUV-HFRS) is characterized by strong neutrophil activation. Neutrophils are the most abundant immune cell type in the circulation and are specially equipped to rapidly respond to infections. They are more heterogenous than previously appreciated, with specific neutrophil subsets recently implicated in inflammation and immunosuppression. Furthermore, neutrophils can be divided based on their density to either low-density granulocytes (LDGs) or “normal density” polymorphonuclear cell (PMN) fractions. In the current study we aimed to identify and characterize the different neutrophil subsets in the circulation of PUUV-HFRS patients. PMNs exhibited an activation of antiviral pathways, while circulating LDGs were increased in frequency following acute PUUV-HFRS. Furthermore, cell surface marker expression analysis revealed that PUUV-associated LDGs are primarily immature and most likely reflect an increased neutrophil production from the bone marrow. Interestingly, both the frequency of LDGs and the presence of a “left shift” in blood associated with the extent of thrombocytopenia, one of the hallmarks of severe HFRS, suggesting that immature neutrophils could play a role in disease pathogenesis. These results imply that elevated circulating LDGs might be a general finding in acute viral infections. However, in contrast to the COVID-19 associated LDGs described previously, the secretome of PUUV LDGs did not show significant immunosuppressive ability, which suggests inherent biological differences in the LDG responses that can be dependent on the causative virus or differing infection kinetics.

## Introduction

Hemorrhagic fever with renal syndrome (HFRS) is caused by Old World orthohantaviruses including Puumala (PUUV) in Northern Europe and Hantaan (HTNV) in Asia. The symptoms of HFRS range from flu-like symptoms to thrombocytopenia, acute kidney injury (AKI), hypotension, hemorrhages, shock and even death (1,2). The exact pathological mechanisms of HFRS are still poorly understood, although many of the clinical findings such as thrombocytopenia and increased coagulopathy can be explained by increased vascular leakage, which in turn is thought to be immunologically mediated, given that the virus itself is unable to cause cytotoxicity in infected endothelial cells (3).

Neutrophilic granulocytes have gained a lot of attention recently as one potential factor driving vascular leakage and kidney dysfunction in HFRS. Recent studies highlight the ability of neutrophils to bind to PUUV-infected endothelial cells and become activated through degranulation and release of neutrophil extracellular traps (NETs) in the acute stage of the disease (4,5). Neutrophils are the most abundant immune cell type in the circulation and play a critical role in the early immune response to infections, exhibiting rapid mobilization and recruitment to sites of inflammation (6,7). In addition to NETs, neutrophils can release various factors, including reactive oxygen species and proteases, which can lead to increased endothelial permeability and vascular leakage (8,9).

In addition to HFRS, neutrophils are strongly involved in the pathogenesis of other acute viral infections such as dengue hemorrhagic fever, COVID-19 and pneumonia caused by respiratory syncytial virus (RSV) or influenza (10). The expansion and activation of neutrophils during acute viral infections is believed to be part of a strong proinflammatory response, especially in severe cases, and there is evidence of increased numbers of immature low-density granulocytes (LDGs) in the circulating blood, a phenomenon we have previously described in the context of COVID-19 (11). This presence is attributed to an emergency myelopoiesis triggered by viral infection and aimed at replenishing the terminally activated neutrophils, thus increasing the number of neutrophils able to counteract virus infections (12).

In the current study we aimed to characterize the neutrophil responses further during PUUV infection by investigating the potential presence of LDGs in acute and subacute HFRS. LDGs are defined by their presence in the peripheral blood monocular cell (PBMC) fraction, as opposed to the denser cellular fractions that include mature granulocytes (polymorphonuclear cells or PMNs) and platelets. Thus, we employed multicolor flow cytometry to identify and quantify LDGs in PBMCs isolated from patients with PUUV-caused HFRS. In addition, we performed transcriptomic and functional analysis of isolated HFRS-associated LDGs and PMNs to better understand their potential role in HFRS disease pathogenesis.

## Materials and methods

### Patient populations

Three different HFRS patient cohorts were included in this study: a Tampere University Hospital (TUH) clinical cohort, a Helsinki University Hospital (HUH) experimental cohort, and a Hantaan (HTNV-HFRS) single cell sequencing cohort.

### TUH cohort

The differential counts from patients with acute PUUV-caused HFRS (confirmed serologically) (13) was performed by routine laboratory methods. All patients were treated at Tampere University Hospital during 1997-2017. All patients gave informed consent to participate, and the study was approved by Ethics Committee of Tampere University Hospital (RO97166, RO99256, RO4180, RO9206, R15007).

### HUH cohort

The study inclusion criterion for PUUV-infected HFRS patients was a positive IgM antibody test done by Hospital District of Helsinki and Uusimaa Laboratory (HUSLAB). The clinical characteristics of the patients (n=25) were obtained through retrospective patient record review (Table **2**). As a control group, age and sex matched healthy donors (HC) were included (n = 9, age range 44-60 with mean 53.11 ± 6.37, male:female 4:5). The study was approved by the Ethics Committee of the Hospital District of Helsinki and Uusimaa (HUS/157/2020). All study participants provided written informed consent.

### HTNV-HFRS cohort

We also reused single cell RNA sequencing data from the GSE161354 Public Gene Expression Omnibus database, which included subjects with Hantaan orthohantavirus-induced HFRS. All the six patients profiled were male (aged 25-52, mean 37.83±10.57), while the two healthy donors consisted of a male and a female (aged 29 and 33 respectively, mean 31±2.83). These samples were collected during the acute phase of HFRS, described as within 10 days after the onset of fever, and disease severity was specified as moderate for three of the patients, and critical for the other three.

### Isolation and culture of neutrophils from human blood

Blood samples from PUUV patients and from HCs were collected in EDTA vacutainer tubes and transported to the laboratory. Peripheral blood mononuclear cells (PBMCs) or polymorphonuclear cells (PMNs) were isolated from whole blood by density gradient centrifugation using Ficoll-Paque Plus (GE Healthcare) or Polymorphprep (Axis-Shield), respectively. Standard procedures were followed, including the use of 2 mM EDTA in PBS as the wash buffer and red blood cell lysis with ACK lysis buffer (Lonza by Thermo Fisher). Subsequently, CD66^+^ granulocytes were isolated from the PBMC and PMN fractions using the CD66abce MicroBead Kit (Miltenyi Biotec, Germany) with an MS column, following the manufacturer’s instructions. The positively selected CD66^+^ LDGs and the isolated PMNs were then washed, counted using a TC20^TM^ Automated Cell Counter (Bio-Rad Laboratories, Inc.) with trypan blue staining for dead cell exclusion, and an aliquot of cells was lysed in Trizol reagent (Thermo Fisher Scientific) and stored at –80°C for later extraction of total RNA and subsequent RNA sequencing (RNAseq) analysis.

After isolation, cells were cultured (1 million cells/ml) for 24 hours in growth medium RPMI-1640 supplemented with 10% inactivated FCS, 100 IU/mL penicillin, 100 μg/mL streptomycin, 2 mM L-glutamine (R10) at 37°C. After incubation, cells were centrifuged, and their supernatant was collected and stored at –80°C for further analysis. Soluble factor stimulation assays were conducted by culturing isolated neutrophils from HC and PUUV-HFRS patients (2 million cells/ml) in RPMI 1640 supplemented with 10% fetal bovine serum (R10) at 37°C. Cells were primed with either LPS (20 ng/ml, Sigma Aldrich) or IFN-I (combination of 2.7*104 IU/ml IFN-α and IFN-β, Immunotools) for 4 h, followed by activation with 2.5 µM nigericin (Invivogen) for an additional 4 h. Finally, cells were centrifuged to collect supernatants, later used for various analyses.

### ELISA detection of calprotectin, MPO, IL-6, IL-8, IL-1β and IL-18

The following ELISA kits were purchased from R&D Systems and used according to the manufacturer’s protocol: human calprotectin/S100A8 DuoSet, human MPO DuoSet, catalog no. DY3174, catalog no. DY4570; human IL-6 DuoSet, catalog no. DY206; human IL-8/CXCL8 DuoSet, catalog no. DY208; human IL-1β DuoSet catalog no. DY201; and human IL-18 DuoSet catalog no. DY318. Supernatants were analyzed in a 1:1000 dilution for MPO and calprotectin, and 1:1 for IL-6, IL-8, IL-1β and IL-18. The concentrations of the respective analytes were determined using the standards supplied by the manufacturer, while the absolute concentrations in supernatant were established by multiplying the determined values by the respective dilution factors.

### T cell activation assay

T cells from freshly isolated PBMCs of three healthy donors were enriched through negative selection, through depletion of non-target cells by anti CD14/ CD15/CD16/CD19/CD34/CD36/CD56/CD123/CD235a biotin-conjugated monoclonal antibodies MicroBeads Kit (Miltenyi Biotec), with the use of a LS column. After isolation, T cells were stained with carboxyfluorescein succinimidyl ester (CFSE) proliferation staining dye (Thermo Fisher) and resuspended in R10. Consecutively, T cells (1 million cells/ml) were activated with Dynabeads Human T-Activator CD3/CD28 for T Cell Expansion and Activation (Thermo Fisher) in culture medium, which consisted of LDG or PMN supernatants and fresh R10 in 1:1 ratio. For experiments, supernatants from individual patient LDGs or PMNs during the acute stage were assayed individually (n = 18 LDG, n = 18 PMN). After 6 days of incubation at 37C (5% CO_2_), the samples were analyzed by flow cytometry for CFSE expression to estimate the number of cell divisions and proliferation index, with the use of the proliferation platform of FlowJo software (v10.0.7, BD).

### Flow cytometry

PBMCs or PMNs were separated from EDTA-anticoagulated whole blood by density gradient centrifugation either with Ficoll-Paque Plus (GE Healthcare) or Polymorphprep (Axis-Shield), respectively, using standard procedures including red blood cell lysis with ACK lysis buffer (Lonza by Thermo Fisher). Before cell staining, 1-2×10^6^ cells were incubated with Fc receptor blocking reagent (BioLegend) for 5 min at RT, followed by a fixable dead cell stain (Invitrogen) and surface staining in the dark for 30 min at RT with the titrated fluorescently labeled anti-human monoclonal antibody cocktail: CD3 FITC (clone OKT3, catalog #: 11-0037-42) and CD66b PE-Cy7 (clone G10F5, catalog #: 25-0666-42) from Thermo Scientific; CD56 FITC (clone MEM-188, catalog #: 21270563) from ImmunoTools; CD19 FITC (clone HIB19, catalog #: 302206), CD33 APC (clone P67.6, catalog #: 366605), LOX-1 PE (clone: 15C4, catalog #: 358603), CD11b APC-Cy7 (clone ICRF44, catalog #: 301341) from BioLegend; CD16 BV421 (clone 3G8, catalog #: 562878), HLA-DR BV786 (clone G46-6, catalog #: 564041), and CD15 BV510 (clone W6D3, catalog #: 563141) from BD Biosciences.

Subsequently, cells were centrifugated at 400 x g for 5 min at 22°C and fixed in 2% paraformaldehyde (PFA) for 15 min RT in the dark. Finally, cells were subjected to flow cytometric analysis with a three-laser (Blue/Red/Violet lasers) 14-color Fortessa LSRII cytometer (BD Biosciences). Data was acquired with BD FACSDiva version 8.0.1 (BD Biosciences) software and further analysis was performed with the FlowJo software v10 (BD Biosciences). A proper instrument daily performance check was ensured by running CS&T beads (BD Biosciences). PBMCs were selected by means of the morphology gate (FSC-A/ SSC-A plot) and gated in the forward scatter by the Height (FSC-H) to Width (FSC-A) ratios to exclude doublets. Only live cells were included in the analysis, and LDGs were defined as CD3^-^/CD56^-^/CD19^-^/CD14^-^/CD66^+^/CD15^+^ cells.

The datasets underwent normalization, followed by concatenation of the samples. Then, the collected data was subjected to unsupervised dimensionality reduction and identification of cell populations, by Uniform Manifold Approximation and Projection for Dimension Reduction (UMAP) (14). To generate UMAP plots, the minimum distance was set at 0.5 and the nearest neighbors’ distance was set at 15, using a Euclidean vector space.

### RNA sequencing

Neutrophils were isolated from four different cohorts: healthy control polymorphonuclear neutrophils (HC PMNs, n = 4), PUUV-infected polymorphonuclear neutrophils (PUUV PMNs, n = 4), PUUV-infected CD16^-^ low-density granulocytes (LDGs) (n = 4, classified as CD16^-^ if containing 80% or more CD16^-^), and PUUV-infected CD16^+^ LDGs (n = 4, comprising the remaining samples after the CD16^-^ selection). After isolation, neutrophils were lysed in Trizol (Thermo Scientific) and the RNA extracted in the liquid phase using chloroform. RNA isolation was carried out using the RNeasy micro kit (Qiagen). Isolated RNA underwent mRNA transcriptome sequencing. Briefly, RNA sequencing was performed using the NEBNext Ultra II directional RNA library preparation method. Sample quality and integrity were assessed using TapeStation RNA analysis. Sequencing was conducted on the Illumina NextSeq platform, followed by standard bioinformatics analysis for gene expression quantification. The service was provided by the Biomedicum Functional Genomics Unit at the Helsinki Institute of Life Science and Biocenter Finland at the University of Helsinki.

### RNA data analysis

Principal Component Analysis (PCA) was performed on the normalized gene expression matrix using the Chipster software (15) to identify patterns in the data and reduce the dimensionality of the dataset. The top principal components were selected based on the percentage of variance explained. Gene Set Enrichment Analysis (GSEA), making use of the KEGG or Reactome databases, was performed on the top differentially expressed genes (DEG) between groups, identified by DESeq2 (adjusted P value < 0.05, log2FC >1) using ExpressAnalyst (16). The resulting p-values were corrected for multiple testing using the Benjamini-Hochberg method, and pathways with a corrected p-value <0.05 were considered significant. Additionally, relevant inflammasome-related genes were identified using GENESHOT (17). To visualize the expression patterns of the DEG, a heatmap was generated using heatmapper.ca (18) for better visualization.

CIBERSORTx, a machine learning algorithm that infers cell type proportions using a reference gene expression matrix of known cell types (19), was used to perform RNA-seq deconvolution on the gene expression data to estimate the abundance of immune cell types in the samples, using Lasalle *et al.* (20) signature matrix which made use of a published whole-blood single-cell dataset (12). This included the main PBMC populations, comprising neutrophils which were subclassified into mature and immature. To note, the less frequent subsets of granulocytes (eosinophils and basophils) are not considered separately in this signature matrix.

### Single cell RNA sequencing analysis

Reanalysis of GSE161354 (21), a shared expression profiling by high throughput sequencing of a single-cell atlas of the peripheral immune response in patients with Hantaan orthohantavirus-caused HFRS, was performed using Chipster software (15).

### Statistical analyses

Statistical analysis was performed using GraphPad Prism 8.3 software (GraphPad Software, San Diego, CA, USA) or SPSS software version 24 (SPSS Inc., Chicago, IL, USA). Statistically significant correlations between parameters were assessed by calculating Spearman’s correlation coefficients, and differences between groups were assessed with Mann-Whitney or Kruskall-Wallis tests, depending on sample distribution and the number of groups analyzed. Multivariate logistic regression analysis was performed to estimate predictors for blood differential count left shift as the outcome. The predictors inlcuded categorial variables smoking status and gender as well as continuos variables age, minimum thrombocyte counts and maximum blood creatinine levels during hospitalization.

## Results

### Blood “left shift” correlates with thrombocytopenia in acute PUUV-HFRS

A “left shift” in routine blood differential counts typically represent increased numbers of immature neutrophils in the circulation and is a common clinical finding in acute PUUV-HFRS. To get initial insight on the role of immature neutrophils in acute HFRS, we investigated clinical records of hospitalized PUUV-HFRS patients (TUH cohort) for the presence of left shift in blood differential counts and its potential association with relevant disease severity markers. We observed a left shift in 68 % of the patients (Table 1, n = 109, and Figure 1A), which significantly associated with lower thrombocyte cell counts (Figure 1B-C) but not with blood creatinine levels, used as a marker of acute kidney injury (AKI). Furthermore, the significant association between left shift and low thrombocyte counts (p= 0.002) remained after multivariate logistic regression analysis, which included patient age, gender, smoking status and maximum creatinine levels during hospitalization (Figure 1C). Thus, these findings link immature neutrophils to thrombocytopenia, but not AKI, in acute PUUV-HFRS.

**Figure 1.**
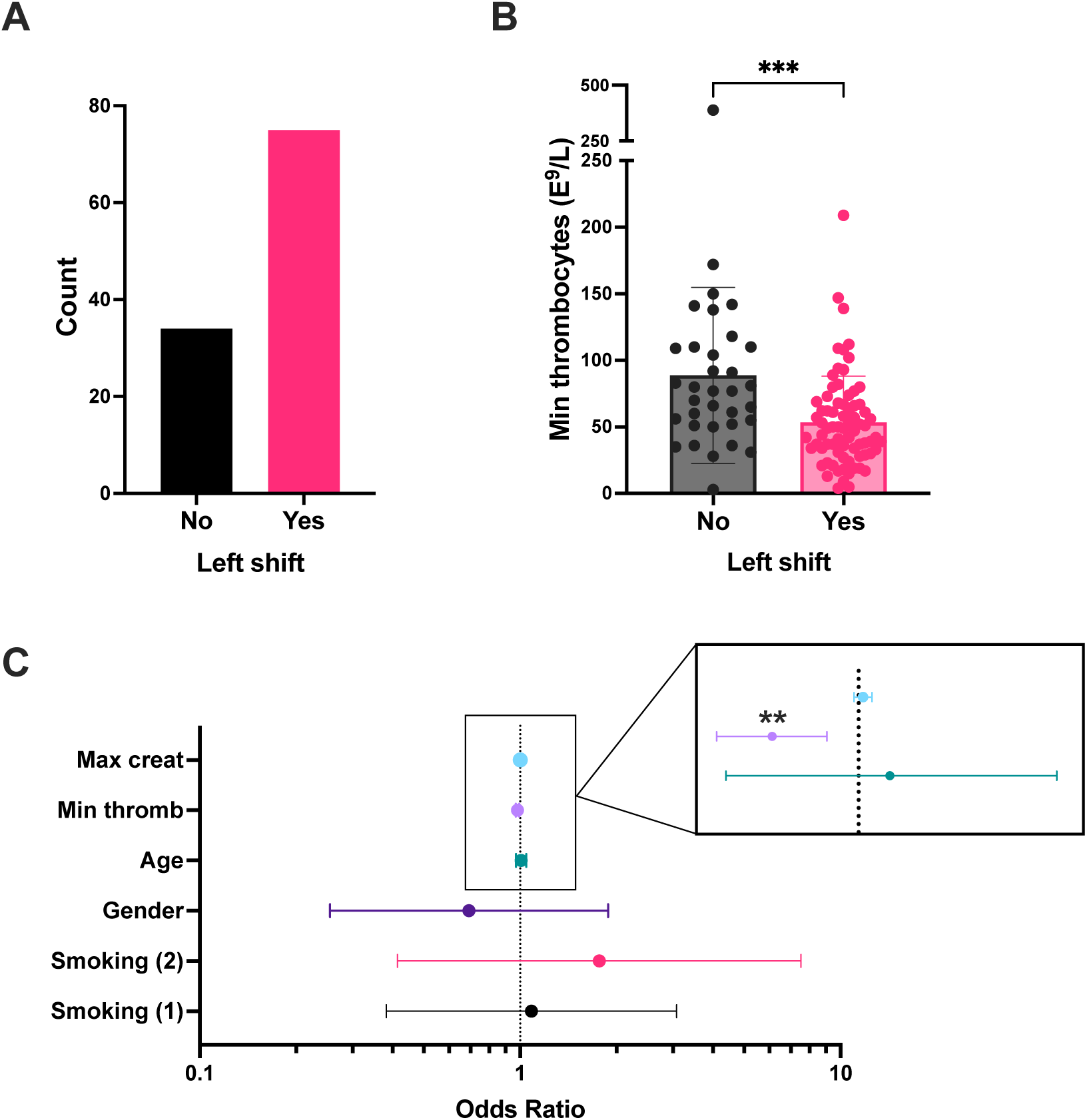
“Left shift” from blood differential counts is linked to thrombocytopenia in PUUV infected patients (TUH cohort). (**A**) The number of PUUV-HFRS patients from the TUH cohort that presented with a “left shift”. (**B**) Minimum thrombocyte counts classified by the presence or absence of differential blood count “left shift”. (**C**) Forest plot depicting the effect of various predictors in a logistic regression model having left shift as an outcome. Each line on the forest plot represents a different predictor variable (e.g., smoking status, age, gender, minimum thrombocyte counts and maximum blood creatinine levels during hospitalization). Minumum thrombocyte counts odds ratio = 0.981, (confidence interval = 0.969-0.993, observed in the enlargement inset) is the only statistically significant predictor. ***p < 0.001 and **p < 0.01. P values for panel B calculated with Mann-Whitney test; and through multivariate logistic regression for panel C. *Max = maximum; creat = creatinine; min = minimum; thromb = thrombocyte counts. Smoking* (*1*)*: current smoker, Smoking* (*2*)*: ex-smoker*

**Table 1.** Characteristics of 109 HFRS patients (TUH cohort). Blood differential counts were obtained from each patient at acute stage of the disease and the presence/absence of left shift recorded. Where applicable, values are indicated by median (minimum-maximum).

| Parameter (unit) | With left shift | No left shift |
| --- | --- | --- |
| Number of patients | 75 | 34 |
| Age (years) | 42.4 (21.5 - 70.6) | 42.9 (21.2 - 67.3) |
| Gender (Male:Female) | 47:28 | 23:11 |
| Days after fever onset (day of sampling) | 5 (1-13) | 5 (2-29) |

### Frequencies of LDGs are increased during PUUV infection

To gain a clearer understanding of the immature neutrophil responses during PUUV-HFRS, we employed flow cytometry to assess LDG frequencies in peripheral blood mononuclear cells (PBMCs) collected from PUUV patients during the subacute or early convalescent phase of infection (HEL cohort, 7-20 days post-infection, n = 23; clinical characteristics of patients shown in Table 2) and in age- and sex-matched healthy controls (HC) (n = 9). We used a combination of features, such as (A) high granularity as indicated by a distinct high side scatter (SSC-A) profile and (B) the cell surface expression of CD15 and CD66b markers, to discriminate LDGs within the PBMC fraction (Figure 2A and B, and Supplementary Figure 1 detailing the gating strategy).

**Figure 2.**
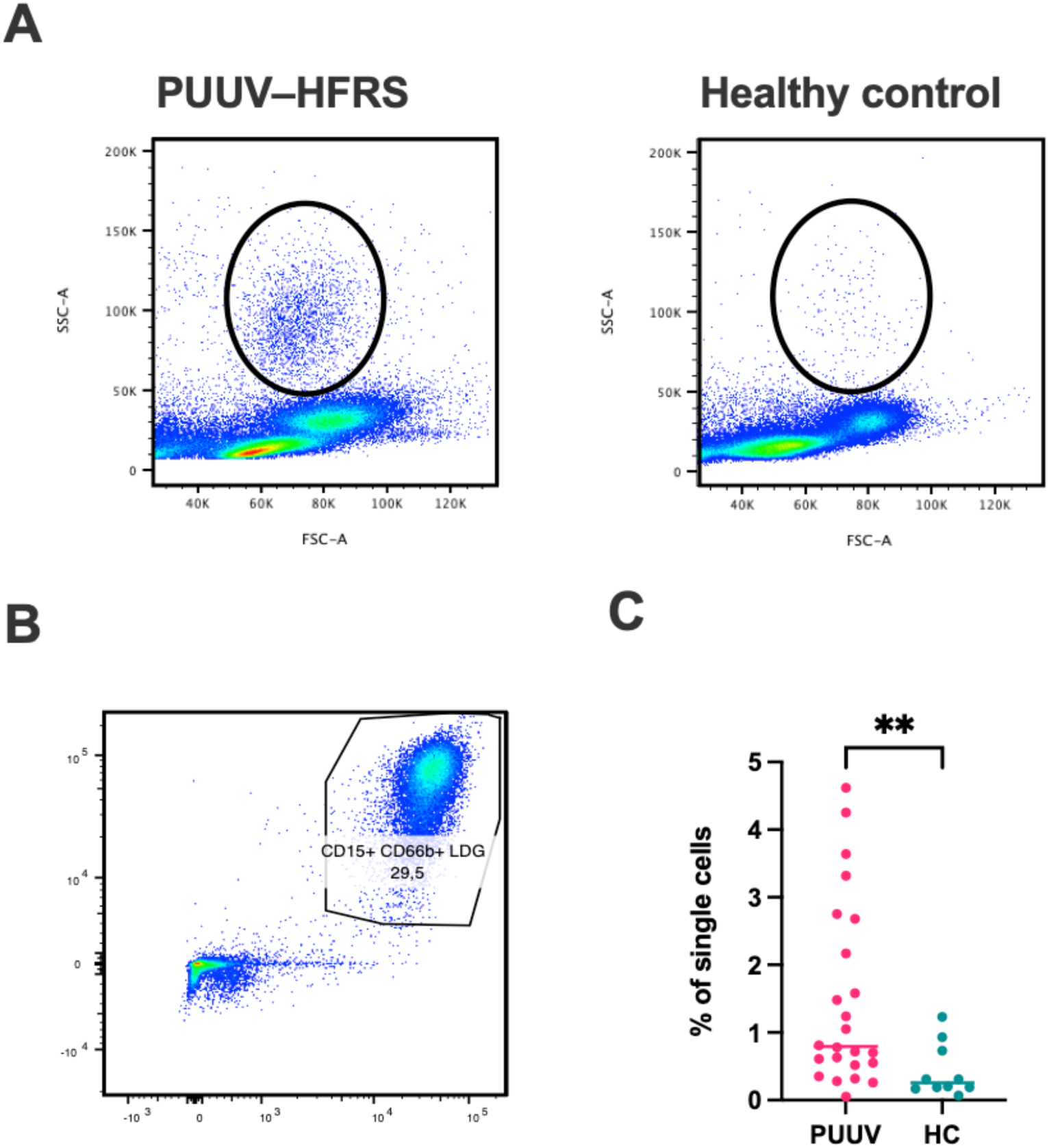
Significantly increased frequencies of LDGs in PUUV-infected patients identified by flow cytometry. PBMCs from PUUV-caused HFRS patients (7-20 days after infection, n = 23) and age and sex-matched healthy controls (HC, n = 9) were isolated from whole blood by density gradient centrifugation and granulocytic cells identified based on their (**A**) high granularity represented by a high side scatter (SSC-A) and (**B**) expression of CD15 and CD66 cell surface markers in single CD56^-^/CD3^-^/CD14^-^/CD19^-^ live cells. (**C**) Quantification of LDG frequencies, calculated as the percentage of single cells.

**Table 2.** Clinical characteristics of 25 HFRS patients (HUH cohort). BP = blood pressure, min = minimum, max = maximum, WBC = white blood cells, ALAT = alanine aminotransferase, CRP = C-reactive protein.

| Parameter (unit) | Range (Mean ± SD) | Reference values |
| --- | --- | --- |
| Age (years) | 35 – 83 ($58.4 \pm 14.9$) | – |
| Gender (Male:Female) | 11:14 | – |
| After fever onset (day of sampling) | 8 – 20 ($13.8 \pm 3.4$) | – |
| Max Neutrophil count (E9/L) | 2.83 – 8.63 ($4.9 \pm 1.6$) | 1.5 – 6.7 |
| Max Systolic BP (mmHg) | 99 – 185 ($134.1 \pm 20.2$) | < 120 |
| Max Diastolic BP (mmHg) | 64 – 102 ($82.7 \pm 10.4$) | < 80 |
| Min Systolic BP (mmHg) | 70 – 143 ($104.7 \pm 18.6$) | < 120 |
| Min Diastolic BP (mmHg) | 47 – 88 ($63.7 \pm 11.7$) | < 80 |
| Max Creatinine ($\mu\text{mol/l}$) | 60 – 571 ($153.6 \pm 131.1$) | Male: 62 – 115 / Female: 53 – 97 |
| Max Hematocrit (%) | 38 – 51 ($43.5 \pm 4.1$) | Male: 39 – 50 / Female: 35 – 46 |
| Min Hematocrit (%) | 29 – 46 ($37.9 \pm 4.2$) | Male: 39 – 50 / Female: 35 – 46 |
| Max Hemoglobin (g/L) | 126 – 180 ($147.7 \pm 15.1$) | Male: 134 – 167 / Female: 117 – 155 |
| Min Hemoglobin (g/L) | 99 – 158 ($128.8 \pm 14.6$) | Male: 134 – 167 / Female: 117 – 155 |
| Max WBC (E9/L) | 4.7 – 18.2 ($8.4 \pm 2.8$) | 3.4 – 8.2 |
| Min Thrombocytes (E9/L) | 26 – 340 ($82.9 \pm 61.8$) | 150 – 360 |
| Max ALAT (U/L) | 19 – 341 ($94.5 \pm 92.3$) | Male: < 50 / Female: < 35 |
| Max CRP (mg/L) | 16 – 197 ($92.9 \pm 47.8$) | < 4 |

We detected significantly increased frequencies of LDGs in PUUV-HFRS patients as compared to HC (Figure 2C), in line with previous reports indicating neutrophil activation during acute PUUV-HFRS (4,5).

Additionally, reanalysis of a single cell RNA-seq data from PBMCs of HFRS patients acutely infected with HTNV (21) revealed an increase in the LDG frequencies, which expanded from 4.3% (3.95% low-density neutrophils and 4.62% low-density basophils) to 8.8% (7.54% low-density neutrophils and 10% low-density basophils) from the total PBMCs isolated from healthy controls vs. HFRS patients (Supplementary Figure 2). This observation confirmed that the increase in LDG frequencies was not exclusive to PUUV-caused HFRS but can be detected also in HTNV-caused HFRS. Furthermore, it suggests that LDGs can be detected already during the acute stages of the disease (6-9 days post onset of fever for HTNV-HFRS patients).

### Expression of the maturation marker CD16 facilitates distinct subdivision within neutrophil populations in PUUV-HFRS

Based on previous observations in other disease contexts, LDGs differ from PMNs by being more immature and showing diminished levels of neutrophil maturity markers CD16 and CD11b (11,22–27). To identify the distinct features of PUUV LDGs in contrast to PMNs, we conducted a phenotypic analysis of neutrophils in PBMC and PMN cell fractions during early convalescent PUUV-HFRS focusing on relevant activation and maturation cell surface markers CD11b, CD16, CD33, LOX-1, and HLA-DR. The UMAP visualization (Figure 2A) provided a consolidated view of single live CD56^-^/CD3^-^/CD19^-^/CD14^-^/CD15^+^/CD66b^+^ cells, effectively highlighting the phenotypic disparities between LDGs and PMNs. Notably, we observed that CD16 expression in PMNs was ubiquitous, with nearly all PMNs exhibiting high CD16 expression (Figure 3B-C), indicative of their maturity. In contrast, CD16 expression in LDGs was more variable, with some LDGs showing varying degrees of CD16 expression while others were negative (Figure 3C). CD11b expression was also higher in PMNs, consistent with their mature phenotype, while LDGs had a higher expression of CD33, representative of their immaturity. Additionally, although remaining at a low level, HLA-DR showed slightly higher levels in PMNs than in LDGs (Figure 3B). Finally, LOX-1 expression was variable and did not significantly depend on the cell type analyzed.

**Figure 3.**
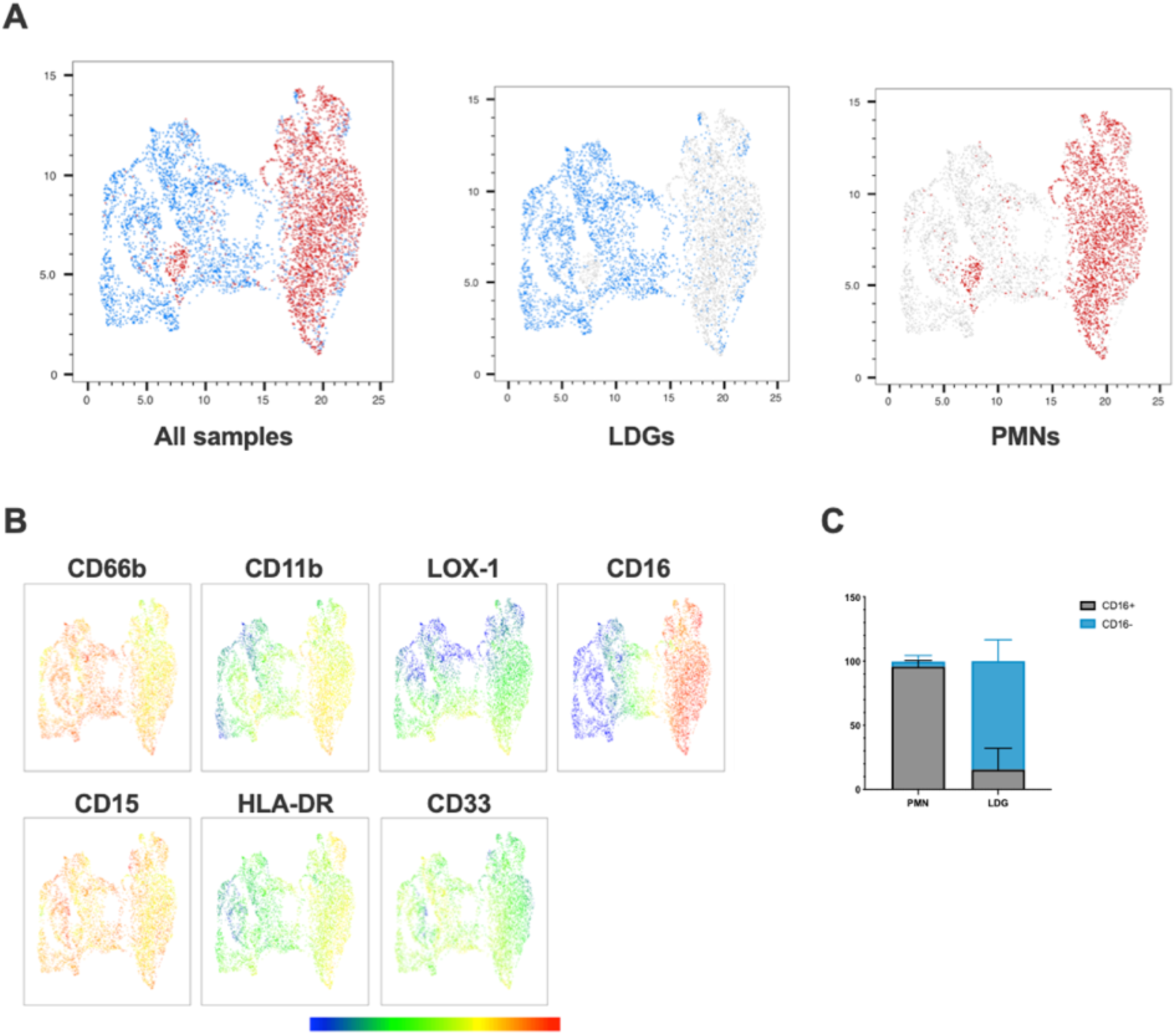
Phenotypic analysis reveals distinct characteristics of PUUV LDGs and PMNs. (**A**) UMAP visualization of single live CD56^-^/CD3^-^/CD19^-^/CD14^-^/CD15^+^/CD66b^+^ cells, displaying LDGs (in blue) and PMNs (in red) in a concatenated format, enabling an in-depth examination of their phenotypic differences. (**B**) Multigraph color mapping depicting each fluorochrome’s Mean Fluorescence Intensity (MFI) for selected markers, including CD66b, CD11b, LOX-1, CD16, CD15, HLA-DR, and CD33. (**C**) Bar graph illustrating the proportion of PMNs and LDGs expressing CD16.

### CD16^+^ LDGs associate with thrombocytopenia in PUUV-HFRS

To assess the clinical significance of the LDG response in PUUV-HFRS, the levels of relevant clinical and laboratory parameters underwent a Spearman correlation analysis (Figure 4A). The clinical parameters were collected at the acute stage of the disease, whereas LDG frequencies were mostly collected at subacute or early convalescent phase. Notably, the analysis revealed a negative correlation between CD16^-^ LDG frequencies with the day after onset of fever at which the sampling took place, indicating potential dynamic changes in the LDG composition over the disease course. As expected, the maximum neutrophil count correlated significantly with the maximum leukocytes from the patients, underlining the connection between overall leukocyte levels and the specific increase in neutrophils during infection. Maximum creatinine levels positively correlated with hospital length of stay, highlighting the link between the duration of hospitalization and reduced renal function, indicative of disease severity in the context of HFRS. Other clinical parameters associated with disease severity include CRP and thrombocytopenia, both of which correlated significantly with the measured LDG frequencies.

**Figure 4.**
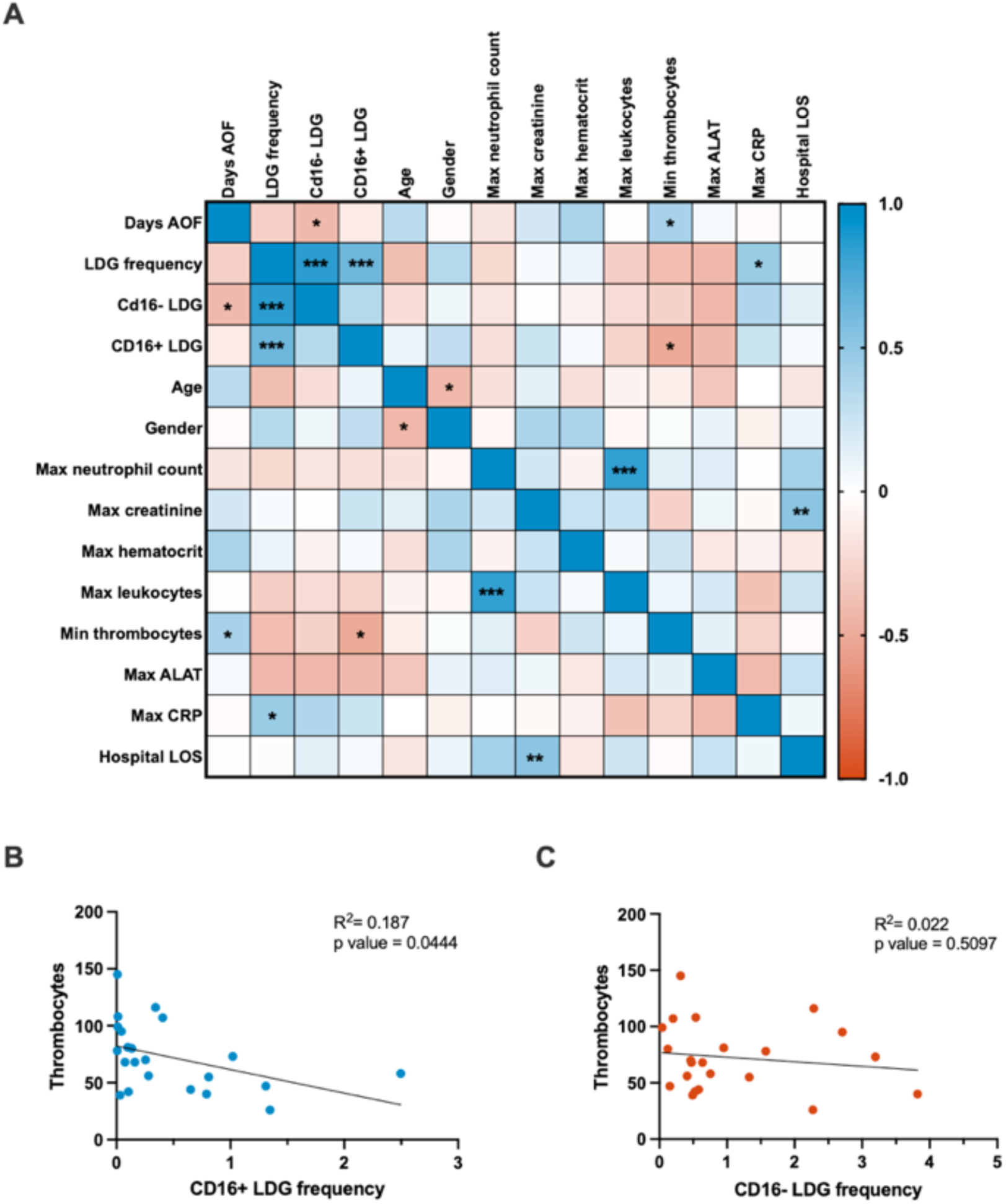
Clinical correlations and regression analyses. (**A**) The correlation matrix presents associations among clinical parameters and flow cytometry-measured expression of CD16 marker in LDGs. Positive and negative correlations are denoted by blue and red colors, respectively. Linear regression analyses were conducted to investigate the relationship between thrombocyte levels and the frequency of (**B**) CD16^+^ LDGs and (**C**) CD16^-^ LDGs, as depicted in scatter plots. The p-value for CD16^+^ LDG regression with thrombocytes was 0.0444, indicating a significant negative association. In contrast, the CD16^-^ LDG regression with the same clinical parameter yielded a p-value of 0.5097, signifying a nonsignificant association with thrombocyte levels. *AOF = after onset of fever; LDG = low-density granulocyte; max = maximum; min = minimum; ALAT = alanine aminotransferase; CRP = C reactive protein; LOS = length of stay*.

Building on these observations, linear regression analysis was employed to delve deeper into the relationship between LDG subsets and thrombocytopenia. This revealed a significant negative association between thrombocyte levels and frequencies of CD16^+^ LDGs (Figure 4B), thereby suggesting a link between the expansion of CD16^+^ LDGs and thrombocytopenia in the context of PUUV-HFRS. On the other hand, the nonsignificant correlation between CD16^-^ LDG frequency and thrombocyte levels (Figure 4C) suggests that this specific LDG subset may not play a significant role in thrombocytopenia development during PUUV infection. The distinct behavior of CD16^-^ LDGs in this correlation analysis highlights the complexity of the immune response during PUUV-HFRS, where different LDG subsets may have variable effects on clinical parameters.

### Transcriptional profiling reveals clear differences between LDGs and PMNs

To gain a comprehensive understanding of the transcriptional differences between neutrophil subsets in response to PUUV infection, we conducted RNA-seq analyses on isolated neutrophil populations. The study included samples from four different groups: healthy control PMNs (HC PMNs, n = 4), PMNs from PUUV patients (PUUV PMNs, n = 4), CD16^-^ LDGs from PUUV patients (n = 4), and CD16^+^ LDGs from PUUV patients (n = 4). The analysis revealed significant differences in gene expression profiles among these groups: principal component analysis (PCA) (Figure 5A) illustrated the separation of these neutrophil populations based on their unique gene expression profiles. Notably, CD16^-^ and CD16^+^ LDGs exhibited a clear segregation from PUUV and HC PMNs, emphasizing pronounced differences between LDGs and PMNs while also showing a clear albeit less pronounced differentiation between PUUV and HC PMNs. Further examination of the differentially expressed (DE) genes between PMNs and LDGs, showcased in a heatmap of the top 150 DE genes (Figure 5B), reaffirms that LDGs possess a unique transcriptional profile when compared to PMNs. Moreover, by estimating the abundance of immune cell types in the samples through RNA-seq deconvolution of the gene expression data, we could confirm that LDGs represented predominantly immature neutrophils and PMNs were primarily mature neutrophils (Figure 5C). However, the CD16^+^ and CD16^-^ LDGs did not differ significantly in this analysis.

**Figure 5.**
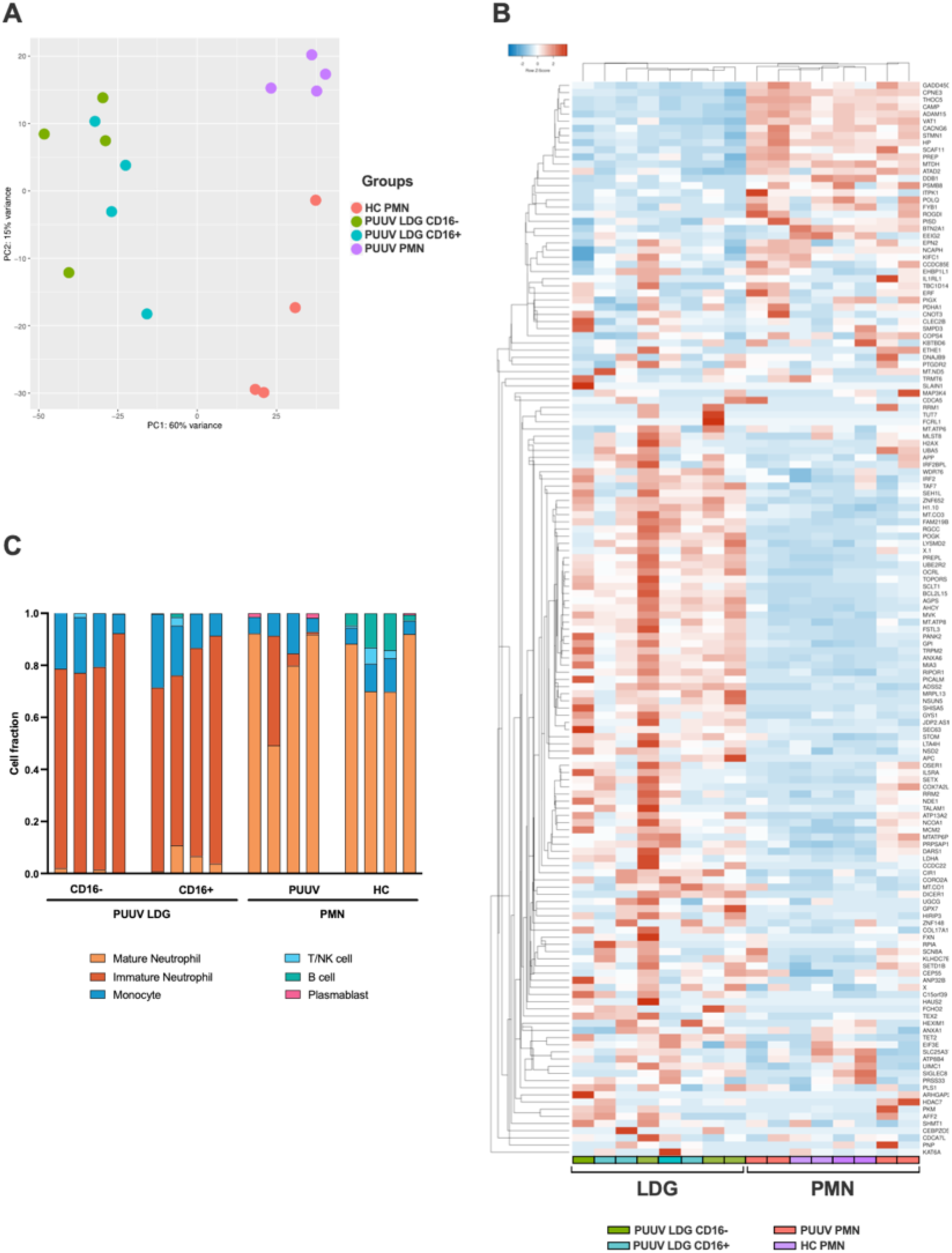
RNA-seq analysis exhibiting the gene expression profile between LDGs and PMNs. (**A**) Principal component analysis (PCA) of the RNA-seq samples (n = 4 for HC PMNs, n = 4 for PUUV PMNs, n = 4 for PUUV CD16^-^ LDGs and n = 4 for PUUV CD16^+^ LDGs). (**B**) Heatmap of the top 150 differentially expressed genes between PUUV PMNs and LDGs, identified by DESeq2. (**C**) Deconvoluted RNA-seq data. The cellular composition in isolated PMN and LDG fractions was estimated using CIBERSORTx through the identification of cell populations based on RNA-seq. The bar plots in the figure represent the cell composition of each RNA-seq sample, offering insights on sample purity (n = 16 samples: 4 PUUV CD16^-^ LDGs, 4 PUUV CD16^+^ LDGs, 4 PUUV PMN and 4 HC PMN).

### Transcriptomic alterations in PUUV patients’ PMNs comprise of inflammasome and interferon-related pathways

To understand the effects of PUUV infection on PMNs, we performed a comparative gene expression analysis specifically between PUUV and HC PMNs. The volcano plot (Figure 6A) reveals distinctive patterns of gene expression, with notable upregulation of interferon (IFN)-related genes in PUUV PMNs. These cells also upregulated genes associated with cell adhesion molecules (CD24, CEACAM6, CEACAM8) during PUUV, which may indicate an increased interaction and communication between immune cells and infected tissues during PUUV, while the upregulation of matrix metalloproteinase-related genes (such as MMP8) may indicate tissue remodeling or repair processes. Additionally, upregulation of genes coding for defensins (DEFA3 and DEFA4), CAMP, ELANE and MPO aligns with an enhanced antimicrobial response during infection, while CEBPE upregulation suggests an active maturation process or response to infection, as this gene codes for a transcription factor involved in granulocyte differentiation (28). Collectively, this gene analysis suggests a coordinated response by PMNs to combat the infection, involving enhanced immune signaling, cell adhesion, differentiation, and antimicrobial activities.

**Figure 6.**
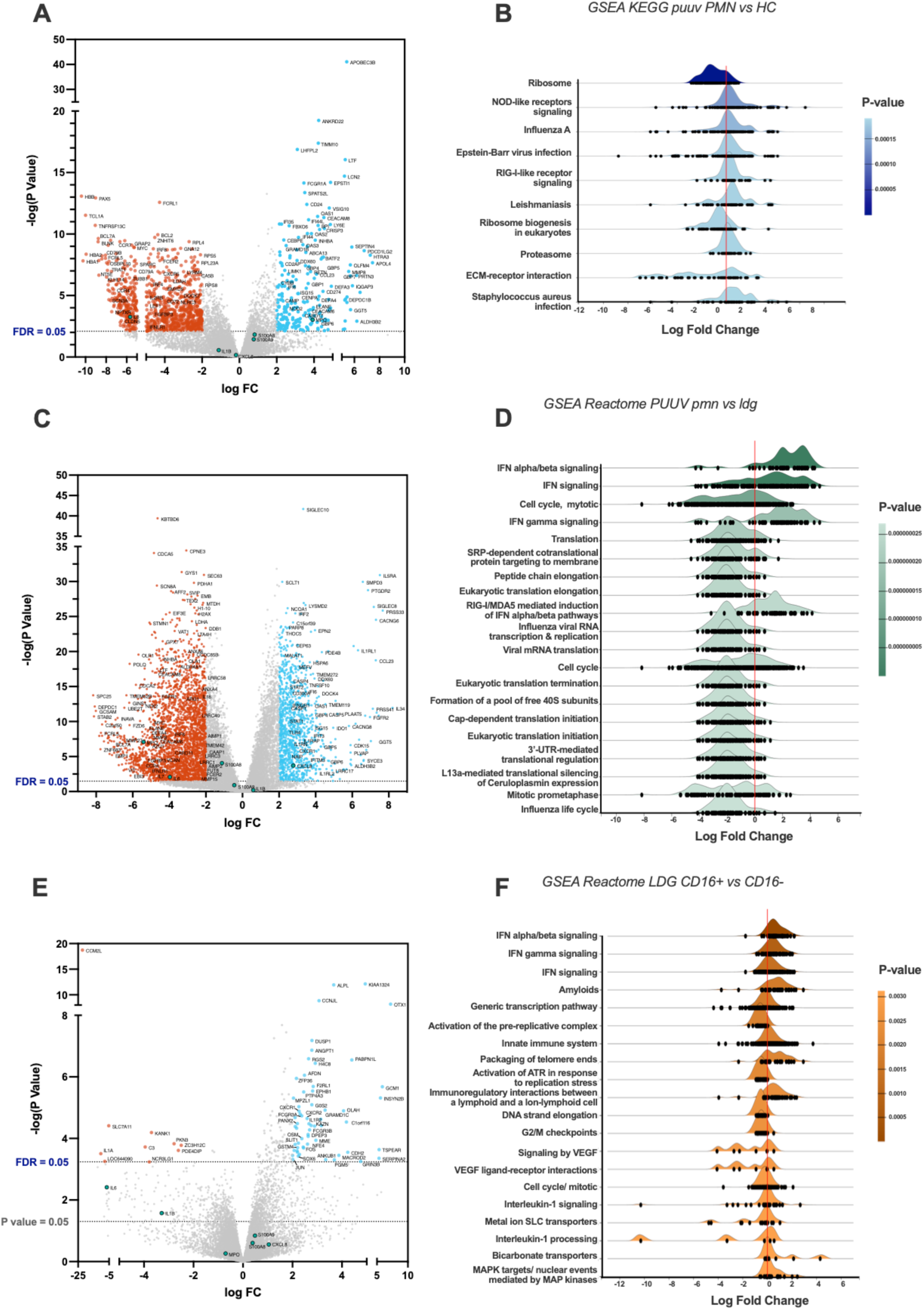
Comparative gene expression analysis between groups. (**A**) Volcano plot highlighting differentially expressed genes (DEGs), with genes upregulated in PMNs during PUUV-HFRS on the right side and downregulated genes on the left, when compared to PMNs from healthy controls (HC). (**B**) Ridgeline diagrams of gene-set enrichment analysis (GSEA) using the KEGG database, depicting the top 10 enriched pathways in PUUV-HFRS PMNs, compared to HC PMNs. (**C**) Volcano plot of DEGs in PUUV-HFRS PMNs, as compared to PUUV-HFRS LDGs. (**D**) Ridgeline diagrams of GSEA using the Reactome database, depicting the top 20 enriched pathways in PUUV-HFRS PMNs, in contrast to PUUV-HFRS LDGs. (**E**) Volcano plot of DEGs in CD16^+^ LDGs, as compared to CD16^-^ LDGs. (**F**) Ridgeline diagrams of GSEA using the Reactome database, depicting the top 20 enriched pathways in CD16^+^ LDGs, in contrast to CD16^-^ LDGs. Genes with a log fold change (FC) greater than 2 and an adjusted p-value (false discovery rate or FDR) < 0.05 are highlighted in the volcano plots, with significant downregulated genes colored in red, and upregulated in light blue. Genes coding for interleukin 6 (IL6) and 8 (CXCL8), myeloperoxidase (MPO), Calprotectin (S100A8 and S100A9), interleukin 1 beta (IL1B) are marked in green, independently of their statistical significance. All GSEA were made using ExpressAnalyst and are sorted by p-value, obtained from Welch’s t-test.

To further understand the biological differences between PUUV-infected and HC PMNs, we employed bioinformatics tools that assess the functional relevance of gene expression data by identifying sets of genes involved in predefined cellular process using gene-set enrichment analysis (GSEA). This analysis revealed positive log fold changes in pathways associated with Nod-like receptor (NLR) and RIG-I-like signaling, proteasome, viral, bacterial and parasitic infections in PUUV-infected PMNs (Figure 6B), indicating an active engagement of responses to infections and immune signaling pathways within PMNs during PUUV infection. On the other hand, ribosome-related pathways were downregulated, suggesting a potential modulation of cellular functions prioritizing immune responses over general protein synthesis.

Furthermore, the upregulation of IFN-related pathways in PMNs from PUUV-infected patients was also evidenced in the DEG analysis against PUUV-infected LDGs, visible in the volcano plot (Figure 6C) and its respective pathway analysis (Figure 6D), indicating a robust antiviral response by PMNs. Conversely, translation-related pathways during PUUV infection were downregulated in PMNs (Figure 6D), as compared to LDGs from PUUV-infected patients, suggesting a heightened activity in protein synthesis processes such as virus-related functions in LDGs. The specific pathways, such as influenza viral RNA transcription and replication, point towards potential interactions with the virus at various stages of its life cycle. This increased activity of pathways involved in protein translation can also suggest continued protein synthesis and active metabolic processes in LDGs, which are well in line with their immature phenotype.

### CD16^+^ LDGs in PUUV infection exhibit a heightened activation of immune responses compared to CD16^-^ LDGs

We performed a comparative gene expression analysis between CD16^+^ and CD16^-^ LDGs during PUUV infection. A volcano plot (Figure 6E) highlights the contrasting gene expression patterns between the two LDG populations. Expanding our analysis, GSEA identified enriched pathways that showcased positive log fold changes in pathways associated with activation of immune responses, including infection, phagosome, cytokine-cytokine receptor interaction, and antigen processing and presentation (Figure 6F). These findings suggest that CD16^+^ LDGs play a role in combatting this viral infection. Simultaneously, negative log fold changes were identified in pathways related to DNA replication and aminoacyl-tRNA biosynthesis.

### Functional characterization of neutrophils in PUUV infections underscores diverse immune responses among PMNs, LDGs, and their CD16^+/-^ subsets

Next, we characterized the different neutrophil populations in PUUV infection by assessing the spontaneous release of various neutrophil activation markers and cytokines in the supernatants of isolated neutrophils during a 24-h culturing period. We also included supernatants of COVID-19 LDGs, which were collected in a similar fashion earlier (11). In the case of calprotectin (Figure 7A), PUUV PMNs exhibited significantly higher levels compared to HC PMNs, COVID-19 LDGs and PUUV LDGs. For myeloperoxidase (MPO) the levels were markedly higher in PUUV and HC PMNs than in COVID-19 or PUUV LDGs (Figure 7B). However, in contrast, when assessing interleukin-6 (IL-6) levels, PUUV and COVID-19 LDGs displayed a statistically significant elevation compared to PUUV and HC PMNs (Figure 7C). The functional characterization of LDGs CD16^+^ and CD16^-^ subsets during PUUV-HFRS (Figure 7F-J) demonstrated that solely IL-6 levels were significantly different, with higher values in CD16^-^ LDGs than CD16^+^ LDGs (Figure 7H). This finding underscores the potential role of IL-6 in PUUV-HFRS and suggests a distinctive immune response mediated by CD16^-^ LDGs.

**Figure 7.**
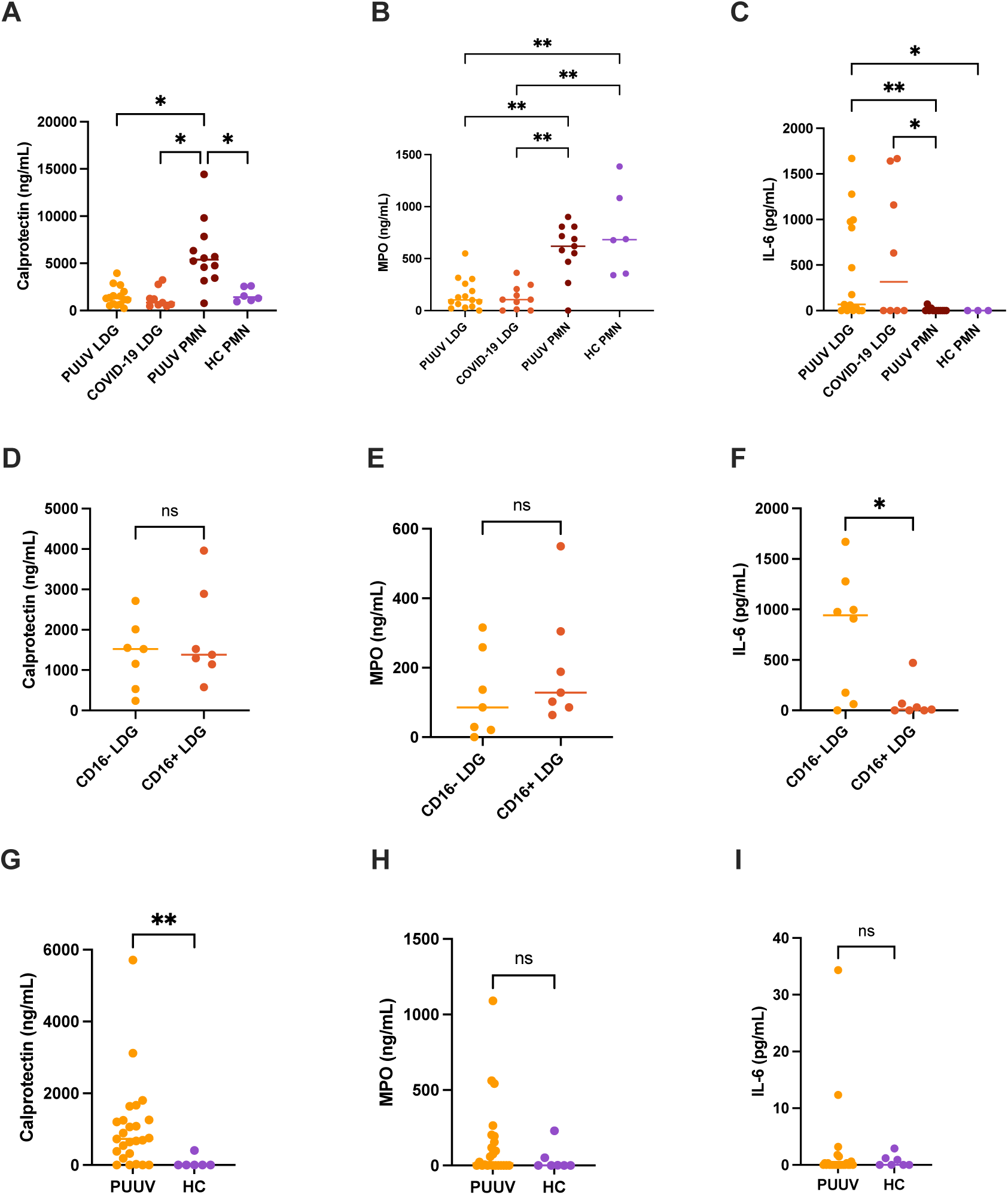
Functional characterization of LDGs and PMNs in PUUV infection. Supernatants of different neutrophil populations cultured for 24-h were collected and analyzed b**y** ELISA. (**A**) Calprotectin, (B) Myeloperoxidase (MPO) and (**C**) Interleukin (IL)-6 levels in PUUV LDGs, PUUV PMNs, HC PMNs and COVID-19 LDGs; and from (D) Calprotectin, (E) MPO and (F) IL-6 levels in CD16^+^ and CD16^-^ LDGs in PUUV-HFRS. (G) Calprotectin, (H) MPO and (I) IL-6 levels were measured from EDTA-plasma samples of PUUV-HFRS patients and healthy controls (HC) by ELISA. *p < 0.05, **p < 0.01, ***p < 0.001. P values were calculated with Kruskall-Wallis test for A-E and Mann-Whitney test for F-J.

These findings prompted us to investigate whether any of these soluble factors could be detected in the circulation at the time of sampling. As expected by the ex vivo cultures, calprotectin levels were elevated in PUUV-HFRS patients in comparison to HCs whereas no significant differences were observed in the case of MPO or IL-6, both of which are known to be increased in acutely hospitalized PUUV-HFRS patients. Taken together, these results highlight the potential intrinsic differences between LDGs and PMNs in their ability to release soluble mediators (IL-6 and MPO, respectively) while PUUV PMNs remain affected by the infection at the time of sampling by specifically releasing calprotectin.

### Suppressed *ex vivo* inflammasome activation in PUUV-HFRS PMNs

Following the detection of upregulated inflammasome-related pathways such as NLR and RIG- I-like signaling in PUUV-HFRS PMNs, we compared the gene expression profile of PMNs and LDGs specifically for several inflammasome-related genes (Supplementary Figure 3). Interestingly, LDGs and PMNs exhibited distinct inflammasome-related gene expression profiles, in which PUUV-HFRS PMNs show the highest expression of some relevant genes, such as CASP1, whereas LDGs were higher in IL-18 gene expression.

To further assess the ability of LDGs and PMNs to engage in inflammasome activation, we explored the release of IL-1β and IL-18 levels upon ex vivo inflammasome priming (type I IFN or LPS) and activation (nigericin) signals in PUUV LDGs (Supplementary Figure 4A and B) and PMNs (Supplementary Figure 4C). Notably, the secretion of both cytokines was increased after *ex vivo* induced inflammasome activation, with PUUV LDGs showing significantly higher levels than PUUV PMNs for both cytokines. For PUUV PMNs, the levels of induced IL-1β remained significantly below HC PMNs, which mean IL-1β responses to both IFN and LPS mediated inflammasome priming were previously observed to be above 200 pg/ml (29). These findings indicate suppression of the inflammasome pathway in PUUV PMNs. which could be due due to previous activation or exhaustion of this pathway during acute PUUV infection. Interestingly, PUUV PMNs showed increased spontaneous release of calprotectin in the 24-h cultures as compared to HC PMNs (Figure 7), suggesting increased activation or cell death of PUUV PMNs that could be linked to their inability to respond to inflammasome activation signals.

### PUUV-HFRS LDGs lack immunosuppressive capacity

Finally, we explored the immunosuppressive capacity of LDGs and PMNs isolated from PUUV-HFRS patients. The PMN secretome of PUUV-HFRS and HCs was able to significantly suppress T cell proliferation as compared to mock-treated T cells. However, LDG secretomes isolated from PUUV-infected patients lacked this ability (Figure 8A). Further LDG classification based on CD16 expression (Figure 8B) highlighted that this behavior was consistent across both groups, irrespective of their CD16 status. These results suggest that the immunosuppressive ability of LDGs is related to disease context and not a general feature of LDGs. However, it remains to be determined whether immunosuppression can be attributed to circulating LDGs present at the acute stage of HFRS.

**Figure 8.**
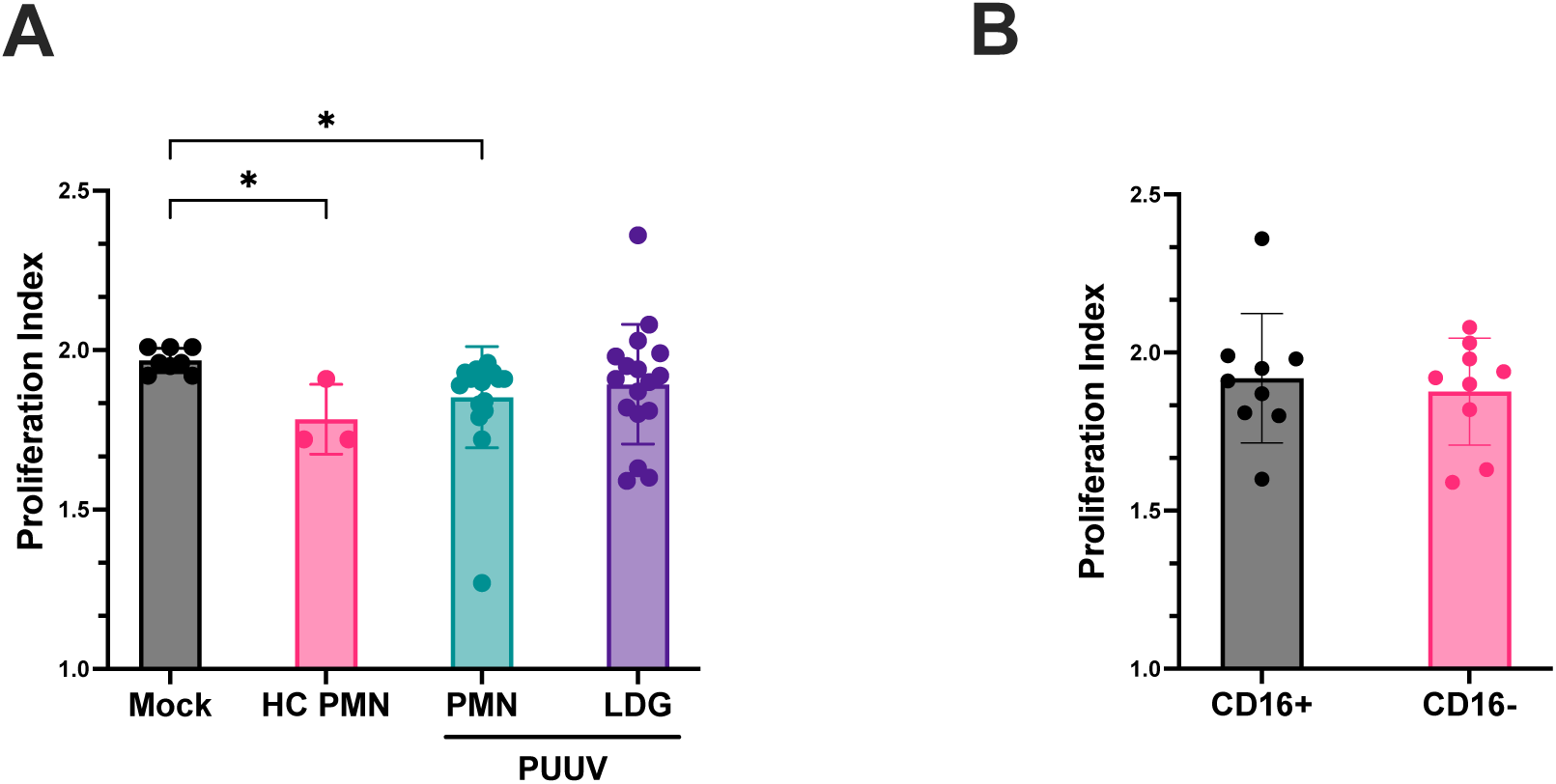
Immunosuppressive function of LDGs and PMNs assay isolated from PUUV-HFRS patients. (**A**) Cell culture supernatants of isolated LDGs and PMNs from PUUV-HFRS or healthy controls (HC) were tested for their ability to suppress T cells in a CFSE-based T cell proliferation assay induced by CD3/CD28 antibodies. The proliferation index was calculated from the total number of divisions divided by the number of cells that went into the division. (**B**) Subclassification of isolated LDG samples based on CD16 expression. *p < 0.05. P values calculated with Kruskall-Wallis test.

## Discussion

The orthohantavirus-caused HFRS is known to be an immunologically mediated disease, but the exact pathological mechanisms remain to be elucidated. The identification of distinct neutrophil populations during PUUV infection enriches our understanding of the intricate immune responses in PUUV-HFRS. Expanding on our prior work on neutrophils during acute COVID-19 (11), we aimed to decipher the phenotypic, functional, and clinical implications of both LDGs and PMNs in the context of PUUV infection. In contrast to mature PMNs, LDGs predominantly exhibited an immature phenotype during acute PUUV-HFRS, suggesting a potential link to increased neutrophil production from the bone marrow (30). Flow cytometry analysis uncovered not only a significant expansion of circulating LDGs during acute PUUV-HFRS, which seems to last at least until early convalescent stage of the disease, but also revealed distinct immunophenotypic profiles within neutrophil populations. Specifically, we identified the cell surface-expressed CD16 as an efficient marker of distinguishing high CD16- expressing mature PMNs from low CD16-expressing immature LDGs. Building upon this, we further classified LDGs into low CD16-expressing (CD16^+^) and non-detectable CD16- expressing (CD16^-^) subsets, shedding light on the functional diversity within the LDG population during PUUV infection.

This immunophenotypic classification was complemented by genotypic studies, employing transcriptomics through RNA-seq analysis, which provided a comprehensive understanding of the underlying molecular signatures driving the observed differences between the different neutrophil populations. When compared to CD16^-^ LDGs, CD16^+^ LDGs displayed an upregulation of pathways associated with immune responses like phagosome, cytokine-cytokine receptor interaction and interferon signaling, whereas CD16^-^ LDGs showed upregulation of genes involved in protein translation and the cell cycle, well in line with their immaturity. The antiviral signaling pathways were further upregulated in mature PMNs, as compared to LDGs in general, and confirmed the immunophenotypic data indicating that CD16^+^ LDGs are further in the differentiation process to become mature neutrophils. These findings also suggest that CD16^+^ LDGs play a role in combatting this viral infectionby increasingly prioritizing immune responses while developing towards fully matured PMNs. The presence of circulating immature neutrophils is typically detected as part of the routine clinical laboratory analysis appearing as a “left shift” of blood differential counts. Indeed, when examining clinical records of hospitalized HFRS patients from the TUH cohort, we identified a left shift in 68 % of the cases. This common clinical finding, attributed to increased immature banded neutrophils, significantly associated with lower thrombocyte cell counts, a key clinical aspect of HFRS severity. Building upon this finding, we also investigated potential links between the increased frequencies of LDG subpopulations and clinical outcomes during the acute stage of PUUV-HFRS patients from the HUH cohort. Intriguingly, the CD16^+^ LDG subset correlated with thrombocytopenia, suggesting a potential link between CD16^+^ LDGs and the severity of HFRS. This finding might be associated with the release of NETs from neutrophils, known to occur during acute PUUV infection (4) The interplay between coagulation and NET formation is well-established and it is documented that NETs can induce platelet aggregation and vice versa (31). Recently, it was observed that thrombocytopenia during PUUV-HFRS could be due to platelet sequestration to infected endothelial cells and/or induced by monocyte-expressed tissue factor (32). Either way, these findings suggest that platelet sequestration or aggregation could be the initial insult to which neutrophils react through NETosis, increased granulopoiesis and finally elevated LDG responses. Conversely, CD16^-^ LDG frequencies were not significantly associated to the extent of thrombocytopenia or other HFRS severity markers.

On the other hand, functional characterization neutrophil subsets in PUUV-HFRS demonstrated higher spontaneous secretion of IL-6 by CD16^-^ LDGs. Given the established association between IL-6 levels and pathogenesis PUUV-HFRS, as indicated by previous studies (33,34), these findings suggest that expansion of CD16^-^ LDGs during PUUV-HFRS could contribute to disease progression by promoting IL-6 mediated proinflammatory signals. On the other hand, IL-6 is known for its ability to promote emergency granulopoiesis through autocrine signaling in hematopoietic progenitor cells (35) and it is possible that the observed spontaneous release in CD16^-^ LDGs could be linked to their differentiation towards mature granulocytes. Taken together, these subtle differences between CD16^+^ and CD16^-^ LDGs offer a refined perspective on the functional diversity within the LDG population during PUUV- HFRS, contributing to the growing understanding of the intricate immune dynamics during hantavirus infections (35,36).

Further comparison of gene expression profiles revealed significant differences between PUUV and HC PMNs, suggesting a potential suppression of ribosomal activities and heightened antiviral responses within PUUV PMNs. Notably, existing research indicates a negative correlation between ribosomal activity and inflammation (37), offering a plausible negative connection between immune response against viral infections and cellular metabolomics. Notable among the increased pathways in PUUV PMNs were NOD-like receptor (NLR) signaling and the proteasome pathway, indicating heightened activity in immune signaling and potential defense mechanisms against viral infections, including inflammasome formation. The activation of inflammasome-related genes and pathways differentiated PUUV PMNs from HC PMNs but also from PUUV LDGs, indicating drastic differences in inflammasome forming potential in neutrophils at different developmental stages.

Inflammasomes are multiprotein structures, which upon stimulation result in the processing and release of pro-inflammatory factors IL-1β and IL-18. Inflammasome formation is well-documented in macrophages but less well known in neutrophils. However, recent evidence by us and others suggests inflammasomes play a major role in neutrophil activation during COVID-19 (29,38,39). To get more insight on inflammasome activation in neutrophils of PUUV-HFRS, we investigated further the inflammasome forming potential of PUUV-HFRS PMNs and LDGs by an ex vivo inflammasome stimulation assay, which involves exogenous priming and activation signals. Surprisingly, we observed significantly diminished IL-1β release in PUUV-HFRS PMNs as compared to PUUV-HFRS LDGs or HC PMNs, suggesting the PMN inflammasome pathway to be exhausted during PUUV-HFRS, possibly due to prior activation in vivo. Accordingly, PUUV-HFRS PMNs also displayed significantly increased spontaneous release of the neutrophil activation marker calprotectin as compared to LDGs and HC PMNs, which may indicate that these cells are in a hyperactivated state and thus less responsive to external stimuli ex vivo. Taken together, these findings suggest alterations in the biology of neutrophils during PUUV-HFRS that are not restricted to the most severe acute stage of the disease but are significantly still present during subacute stages during which recovery from other clinically relevant symptoms is rapidly advancing.

Immunosuppression has been increasingly attributed to LDG populations in cancer and some acute bacterial and viral diseases such as COVID-19 (11,26,40). Examining the immunosuppressive capacity of LDGs and PMNs secretomes isolated from PUUV-HFRS patients revealed a marked contrast. While PMNs, both from PUUV-HFRS and healthy controls, exhibited significant suppression of T cell proliferation, LDGs isolated from PUUV- infected patients lacked this immunosuppressive ability. These findings underscore a clear deviation from the immunosuppressive traits observed in COVID-19 associated LDGs, emphasizing disease-specific variations in LDG functionality. Of note, the COVID-19 LDGs displayed higher levels of CD16 expression than their PUUV-HFRS counterparts in this study that was comparable to mature PMNs (11). Thus, it is possible that the high CD16 expressing LDGs are the ones responsible for the observed immunosuppression during acute COVID-19. However, further experiments using LDGs isolated from early acute HFRS and/or subacute stages of COVID-19 would be required to fully understand these differences.

In conclusion, the expansion of LDGs during acute PUUV infection highlights the importance of investigating the role of neutrophils, including LDGs, in the immune response to hantavirus infections. The potential link between LDGs and clinical outcomes, namely thrombocytopenia, represents an exciting area for further exploration. This study provides the foundation for future investigations aiming to elucidate the functional characteristics of LDGs, their involvement in vascular leakage, and their potential as therapeutic targets in PUUV infection.

## Contributors

L.E.C.: conceptualization, data curation, formal analysis, investigation, software, validation, visualization, writing– original draft; J.T.: investigation, formal analysis; S.T.J.: investigation, writing – review & editing; S.M.: resources; A.V.: resources; writing – review & editing; J.M.: resources, O.V.: funding acquisition, resources;, M.K.: resources; T.S.: conceptualization, data curation, formal analysis, funding acquisition, investigation, methodology, project administration, supervision, validation, writing – original draft.

## Acknowledgements

This work was financed by grants by the Academy of Finland to T.S. (321809); grants by the Helsinki University Hospital funds to O.V. (TYH 2021343); EU Horizon 2020 programme VEO (874735) to O.V.; Paulon Säätiö to L.E.C.; Suomen Lääketieteen Säätiö to L.E.C.; Jane and Aatos Erkko foundation to O.V. The funders had no role in study design, data collection and analysis, nor decision to publish, or preparation of the manuscript.

RNA isolation, library preparations and RNA sequencing was performed at the Institute for Molecular Medicine Finland FIMM, Genomics unit supported by HiLIFE and Biocenter Finland. The authors also thank S. Mäki and M. Utriainen for expert technical assistance.

---

**Supplementary Figure 1.**
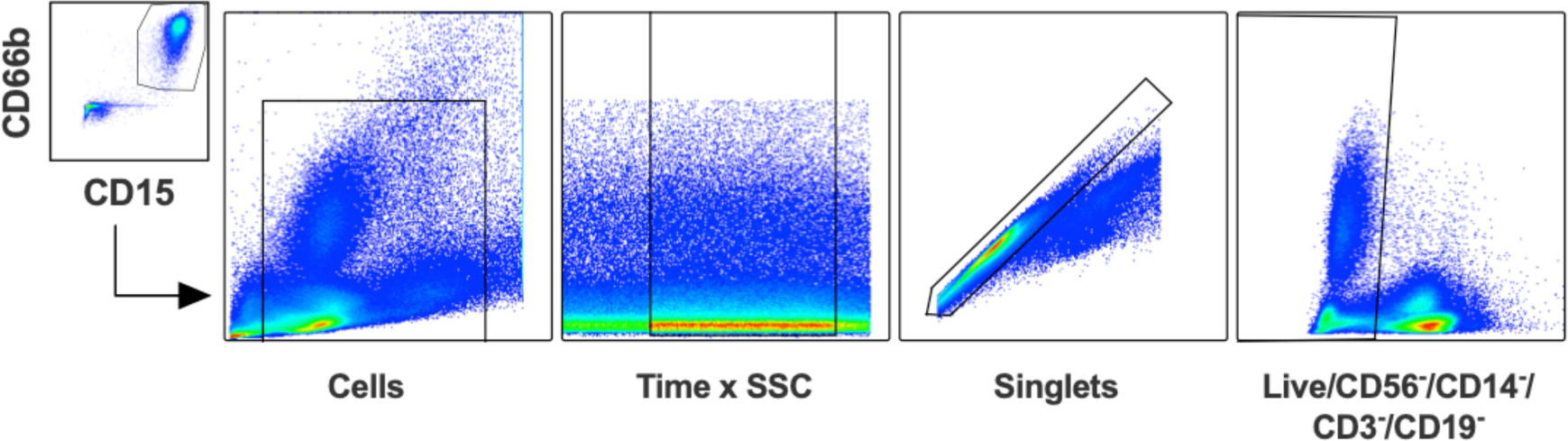
Flow cytometry gating strategy. Single cells were gated after the exclusion of debris and a time gate, which was employed for the selection of a steady collection flow to ensure proper sample quality and homogeneity. Then, cells negative to CD3, CD56, CD14, CD19 and the dead cell marker were gated in, from which CD66b/CD15 double positive cells were finally labeled as LDGs.

**Supplementary Figure 2.**
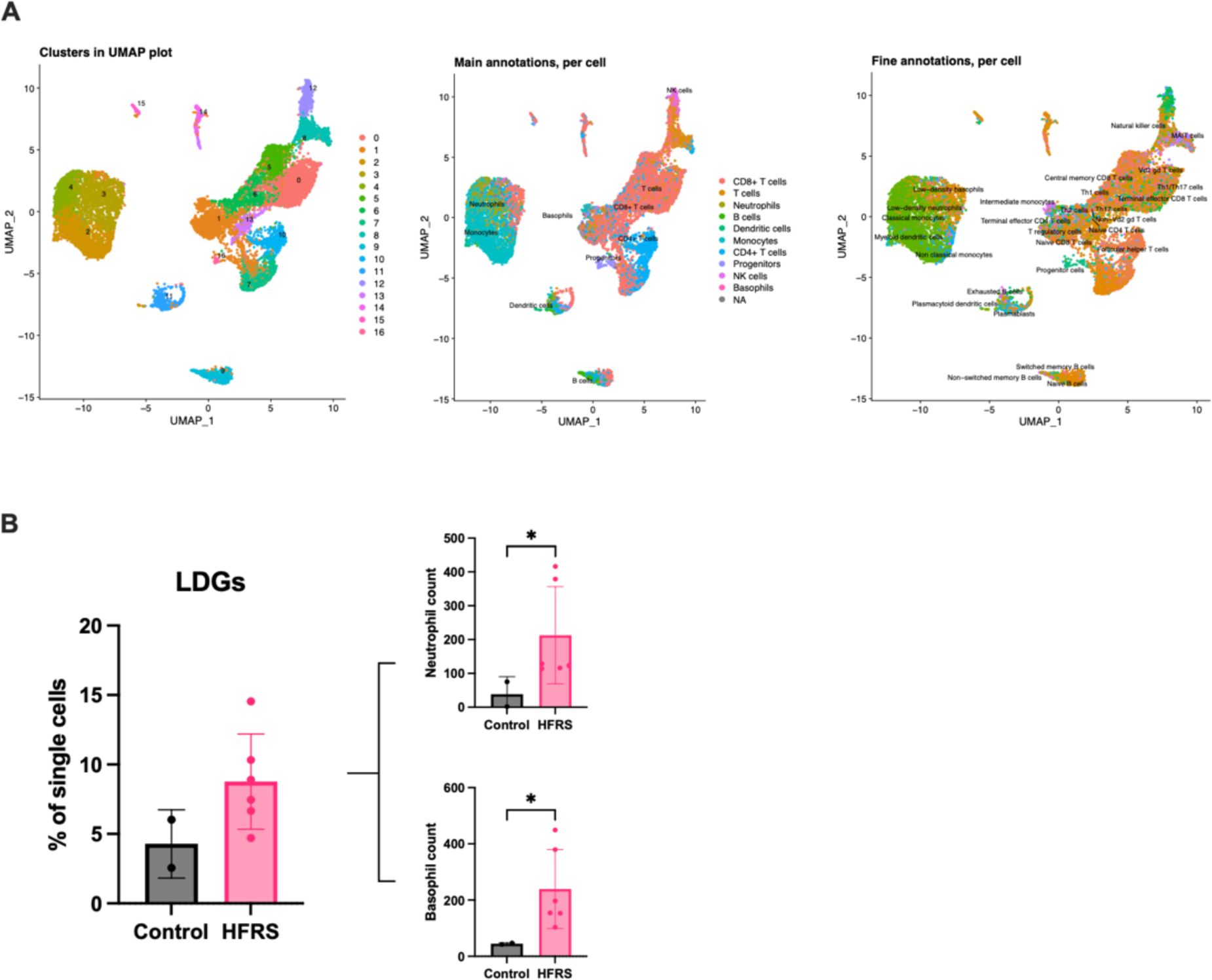
Increase of LDGs in Hantaan orthohantavirus-caused HFRS identified by single cell RNA sequencing analysis. (**A**) UMAP visualization panels depicting the identified numbered clusters. The main annotations are labeled in the UMAP plot using SingleR in Chipster, and the fine annotations are presented, providing additional insights with more clusters and labels. (**B**) Bar plot representing the percentage of LDGs among the total single cells identified by single cell RNA sequencing analysis (scRNA-seq) in both control and HFRS groups. Two smaller bar plots depict the counts of single cells classified as low-density neutrophils and low-density basophils.

**Supplementary Figure 3.**
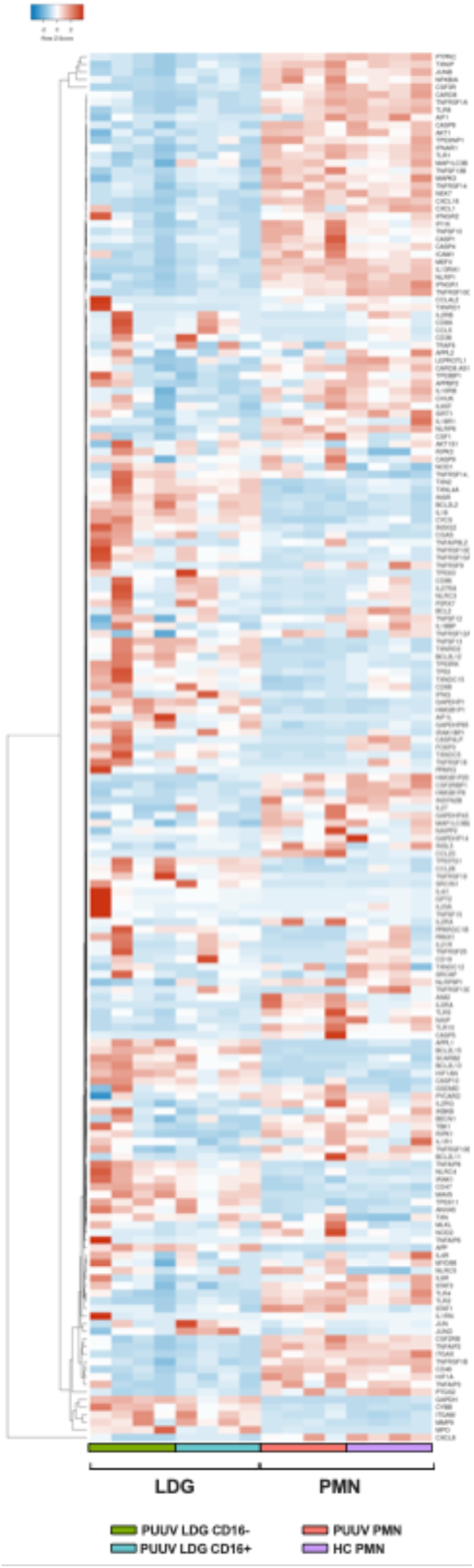
Gene expression of inflammasome-related genes in neutrophils during acute PUUV-HFRS. Heatmap of differentially expressed inflammasome-related genes between LDGs and PMNs.

**Supplementary Figure 4.**
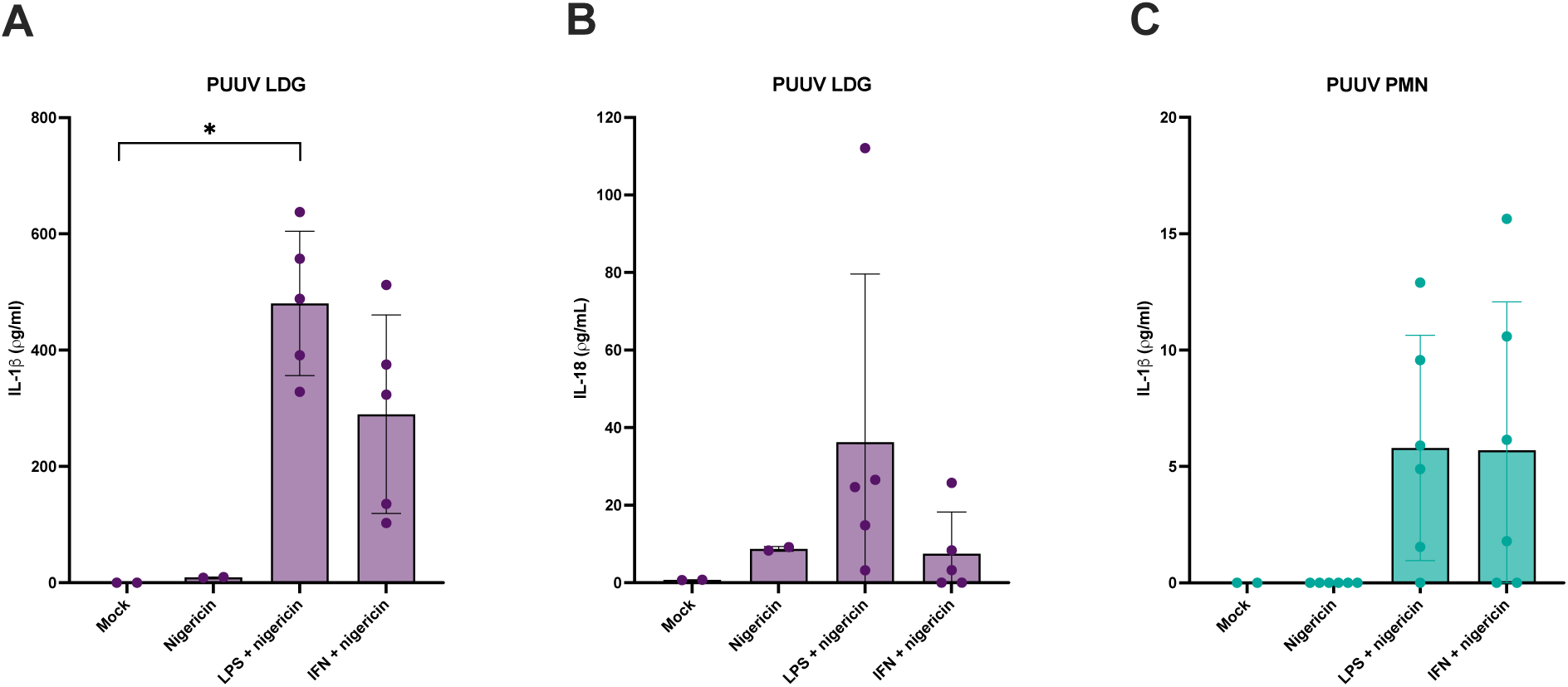
Inflammasome activation in PUUV-HFRS neutrophils. (**A**) IL-1β and (**B**) IL-18 levels in PUUV LDGs, and (**C**) IL-1β levels in PUUV PMNs following interferon or LPS priming and nigericin activation. *p < 0.05. P values calculated with Kruskall-Wallis test.

